# Microbial dysbiosis is evident in oral squamous cell carcinoma (OSCC) tissues of a group of Sri Lankan male patients with oral risk habits

**DOI:** 10.1101/2024.11.19.624363

**Authors:** Manosha Lakmali Perera, Irosha Rukmali Perera

**Affiliations:** School of Dentistry and Oral Health, Griffith University, Queensland, Australia (Alumni); Preventive Oral Health Unit, The National Dental Hospital (Teaching), Colombo, Sri Lanka

## Abstract

**Background:** Metagenomic investigations into components of the oral microbiome in a single study remain scarce.

**Objective:** To ascertain the microbiome profile of oral squamous cell carcinoma (OSCC) tissues in a group of Sri Lankan male patients.

**Methods:** From the main sample of an unmatched case-control study, a sub-sample consisting of 29 OSCC cases and 25 Fibroepithelial polyp (FEP) controls were selected. OSCC incisional and FEP excisional biopsies were collected and stored at -80^0^C. DNA was extracted from frozen specimens using Gentra Puregene Tissue kit (Qiagen, Germany), solid tissue protocol. The DNA extracts were stored at -80^0^C. Extracted DNA samples were sequenced by Illumina’s 2×300 bp chemistry. High quality non-chimeric merged reads were classified to the species level by prioritized BLASTN-algorithm for bacteriome and BLASTN-algorithm with UNITE’s named species sequences as reference for mycobiome.

**Results:** Our study identifies several potential periodontal pathogens, including *Campylobacter concisus, Prevotella salivae, Prevotella loeschii*, and *Fusobacterium oral taxon 204*. Additionally, *Candida albicans* is frequently associated with oral potentially malignant disorders (OPMDs) and oral cancers. Notably, we found a significant association between *Candida etchellii* and oral cancer for the first time. Furthermore, Capnocytophaga, Atopobium, and Candida were the most abundant microbial genera present in oral squamous cell carcinoma (OSCC) tissues.

**Conclusions:** A dysbiotic microbiome was discovered in the tumor microenvironment of a group of OSCC male patients with oral risk habits. Validation of these biomarkers indifferent cohorts to supplement inadequacy in the diagnosis of epithelial dysplasia histologically is much warranted in the golden era of *microbiome-first-medicine*

## 1. Introduction

Polymicrobial infections due to microbial dysbiosis are reported to be responsible for 15 % of infections in immuno-compromised patients with cancers, with no exception with oral cancer [1]. The latest GLOBOSCAN 2018 by the International Agency for Research on Cancer (IARC) featured lip and oral cancer as the 17 ^th^ most common cancer across the globe [2], more prevalent in South Asia and the Pacific Islands [2]. Alarmingly, this cancer type remains 1^st^ and the leading cause of cancer-related death among Sri Lankan males nationally [3, 4]. Moreover, Sri Lanka overwhelms the 5^th^ highest Age Standard Rate (ASR) of incidence of lip and oral cavity cancers globally, with an estimated 2,152 new cases accounting for 9.1% of all types of cancers [2]. Oral squamous cell carcinoma (OSCC) accounts for 90-95% of oral malignancies in most countries (3, 4). Oral cancer consists of malignant neoplasms, evolving from the mucous membranes lining [International Classification of Disease tenth revision (ICD-10)]: Lips (C00); base of the tongue (C01), oral cavity including the anterior two-thirds of the tongue (C02–C06) respectively belongs to the head and neck squamous cell carcinomas (HNSCC) s (5, 6). There are various geographic and population-specific risk factors that contribute both individually and together to the development of oral cancer. Among these, well-established oral risk habits for oral squamous cell carcinoma (OSSC) include smoking, the use of smokeless tobacco, chewing areca nut and betel quid, consuming alcohol, and infections with human papillomavirus (HPV).

The genome of humans inherited from both parents is <150 folds [7] of the acquired metagenome as humans are called ’super-organisms” or “holobionts” with the indigenous microflora [7] mainly comprising bacteriome and mycobiome specific to each individual may demonstrate microbial shifts [8, 9] in metabolic, autoimmune, congenital and neoplastic and other degenerative diseases [8] as well as sleep disorders associated with chronic inflammation and immunosuppression. With the advent of next-generation sequencing (NGS) technologies to view the microbial ecology in unprecedented breadth and depth [7], the human microbiome is rewarding, elevating scrutiny of the role of microbial consortium in non-infectious diseases. Risk panels and microbial signatures related to different cancer types oral, pancreatic, colon, and breast cancer gained the attention of researchers, especially in the past decade. The resulting dysbiotic community progresses the carcinogenesis with their carcinogenic metabolites and toxins. Their virulence factors, cellular components and proteins target the specific aspects of the host’s immune surveillance and immune elimination of neoplastic cells. Consequently, there should be the most eligible bacterial candidates for in vitro and invivo experiments, to view interaction of specific bacteria with the host’s neoplastic cells and immune cells in the tumor microenvironment. In the shed of light, the objective of this study was to use NGS technologies coupled with a species-level taxonomy alignment algorithm to compare the microbiome profile within OSSC tissues to benign intra-oral fibro-epithelial polyps (FEP).

## 2. Materials and Methods

### 2.1 Study design, sample size calculation, setting, subjects and ethical approval

In the present study, a subset was selected representing the vast majority of OSCC cases from an unmatched large case-control study based on the sample size calculation Kelsey JL, Whitmore AS, Evans AS and Thompson WD as published elsewhere [8], of 134 cases and 134 controls as described previously [4, 9, 10, 11].

The subjects were recruited between 17 April 2015 and 2 August 2015 at nine oral and maxillofacial (OMF) units in six provinces of Sri Lanka. Cases comprised 25 Sinhala, ≥40-year-old males with histologically confirmed OSCC affecting the buccal mucosa or tongue. The control group consisted of 27 Sinhala males with a clinical diagnosis of FEP also involving the buccal mucosa or tongue. Subjects with a history of antibiotic use in the last 2 months were excluded.

Ethical approval was granted from the Faculty Research Committee, Faculty of Dental Sciences, University of Peradeniya, Sri Lanka (FRC/ FDS/UOP/E/2014/32) and Griffith University Human Research Ethics Committee, Australia (DOH/18/14/ HREC). Written informed consent was obtained from each participant.

### 2.2 Date collection and clinical oral examination

A pretested, interviewer administered questionnaire was used to collect information on socio-demographics, clinical and oral risk habits, including use of smokeless tobacco, areca nut and betel quid chewing, tobacco smoking, and alcohol consumption. Clinical oral examinations were conducted by dental public-health specialists. The oral mucosa was meticulously inspected for any growth, ulceration, or white patches. The number of missing teeth was recorded. Oral hygiene status was assessed with the simplified oral hygiene index [12], while periodontal status was assessed using bleeding on probing (BOP), periodontal pocket depth (PPD), and clinical attachment loss (CAL) at four sites per anterior tooth and six sites per posterior tooth.

### 2.3 Tissue sampling and DNA extraction

Tissue samples were obtained from incisional biopsies performed to histologically confirm clinically suspected oral squamous cell carcinoma (OSCC) lesions. The freshly collected biopsy was placed on a stack of sterile gauze, and a small piece of tissue (approximately 3 mm^3^) was excised from the deep tissue at the macroscopically visible advancing front of the neoplasm, taking care to avoid contamination from the tumor surface. A new sterile surgical blade was used for each case. The sample was aseptically transferred into a screw-cap vial and placed in a polystyrene container filled with dry ice. These samples were then transferred to a -80°C freezer in a university laboratory as soon as possible. The rest of the biopsy in 10% buffered formalin was sent to the histopathology laboratory. The samples were taken from histopathologically confirmed biopsies were included in the study. Control tissue samples from freshly excised, clinically diagnosed fibroepithelial polyps (FEPs) were collected as described previously [10].

Tissue samples (∼100 mg each) were finely chopped using a sterile blade. DNA extraction was then performed using Gentra Puregene Tissue kit (Qiagen, Hilden, Germany), according to the manufacturer’s instructions (solid tissue protocol) with a few modifications: (1) incubation in the lysis buffer was performed overnight; and (2) an additional lysis step using 50 units of mutanolysin at 37°C for 1.5 h to digest the cell wall of Gram + bacteria was included prior to the addition of Proteinase K (the samples had also been planned to be analyzed for bacterial content). Total DNA concentration and purity were determined using the Nano Drop™ 1000 Spectrophotometer (Thermo Fisher Scientific, Waltham, MA). The extracts were stored at –80°C as published elsewhere [10].

### 2.4 Nucleotide sequencing and taxonomy alignment of bacteriome up to species level

DNA extracted from fresh biopsies was subjected to amplicon library preparation, indexing, and sequencing for the V1 to V3 region with Illumina’s 2 × 300–bp chemistry. High-quality nonchimeric merged reads were classified to the species level with a prioritized BLASTN-based algorithm. Downstream compositional analysis was performed with QIIME (Quantitative Insights into Microbial Ecology) and linear discriminant analysis effect size as described previously [4, 9].

### 2.5 Nucleotide sequencing and taxonomy alignment of mycobiome up to species level

DNA was extracted and an amplicon library was prepared, as previously described [10]. The fungal internal transcribed spacer 2 regions were sequenced using Illumina™ 2 × 300 bp chemistry. The merged reads were classified at the species level using the BLASTN algorithm, referencing UNITE’s named species sequences. Subsequent analyses were conducted using QIIME™ and linear discriminant analysis effect size [10].

### 2.6 Sociodemographic and clinical data analysis

These data were entered and analyzed using SPSS-21 Statistical Package. Percentage distributions were presented as descriptive statistics and Chi-Square Test of Statistical Significance was used as inferential statistics to compare groups with regard to differences in risk habit profiles, oral hygiene status and periodontal disease status. Furthermore, t-test and Fisher’s exact test to compare groups (cell counts <5) were used to compare means with regard to duration of risk habits, missing teeth, mobile teeth, decayed and filled teeth among cases and controls as published [4].

## 3. Results

### 3.1 NGS data processing of profile bacteriome

Seven samples (4 OSCC cases and 2 FEP controls) were excluded due to poor quality read counts or primer mismatches and were excluded. The sequencing run has generated total of 3,277,451 raw paired reads in all, 53 samples. After removal of primer mismatches, additional reads filtering followed by alignment and chimera checking, a total of final 451,048 (40.27%). High-quality, non-chimeric merged reads were obtained with an average length of 482.5 bp. OF these 65 reads were identified de novo as additional chimeras; 61 did not return BLASTN matches and 47,038 formed singleton OTUs and were thus excluded. The number of classified reads per sample ranged from 4358 to 21251 reads. The means (± SD) read counts of the bacteriome were 9102.79 (±3245. 63). Finally, 394,937(35.26%) of the merged reads of 25 OSCC cases and 22 FEP controls were assigned into species level

### 3.2 NGS data processing of mycobiome profile

Seven samples were ended up with low read counts (<3000) or with very high count (an outlier).The sequencing run generated 1,576,427 raw paired reads. About 14% of these were discarded due to primer mismatches; 96% of the remaining reads were successfully stitched with PEAR. Quality filtration and chimera checking removed around 8% of the merged reads leaving a final of1,063,430 (67.5%) reads, 205-535 bp long. Around 95.7% of these reads from 22 OSCC cases and 25 FEP controls were successfully classified to the species level; 2.3% did not return BLASTN matches and 2% formed singleton OTUs and were thus excluded. The number of classified reads per sample ranged from 3973 to 54849 reads (mean of 21,641±13,942).

### 3.3 Overall bacteriome compositional profile

A total of 1,072 species-level taxa (including 373 potentially novel species) belonging to 272 genera and 19 phyla were detected in the 48 samples as scientifically published [4, 9].

### 3.4 Bacterial species richness and diversity of OSCC vs FEP

The number of species identified in the cases and controls was 688 and 810, respectively, with 426 species in common. The number of species ranged from 20 to 183 in the OSCC group and 23 to 229 in the FEP control group samples. Species richness and α-diversity were higher in the controls as compared with the cases, but the differences were not statistically different (Table 1). In PCoA, the cases and controls tended to form separate clusters based on binary data (unweighted UniFrac) and abundance data with the clusters as seen in Figure1, being statistically different as assessed by analysis of similarities (*P* = 0.01 and 0.03, respectively) as scientifically published [4, 9].

**Figure 1:**
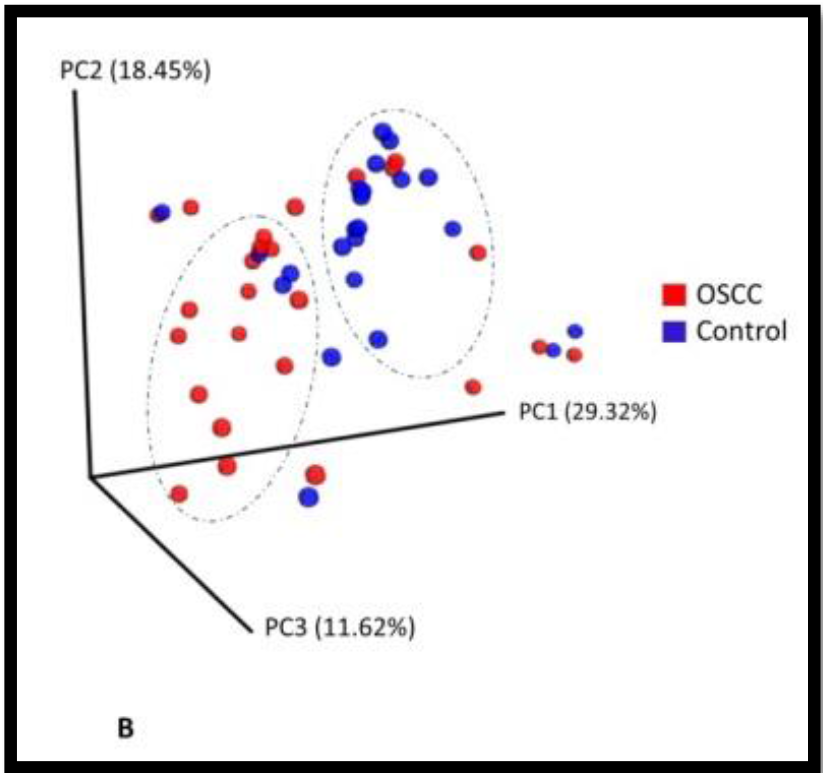
β-diversity analysis. Clusters formed by principle coordinate analysis (PCoA) (weighted Unifrac).

**Table 1:** Species Richness, α-diversity, and Coverage Calculated from the Rarefied BIOM.

| Group | Observed richness | Chao1 | Shannon index | Good's coverage |
| --- | --- | --- | --- | --- |
| OSCC | 82.9±37.8 | 111.2±51.5 | 3.3±0.9 | 0.994±0.003 |
| FEP | 101.6±42.9 | 125.2±50.9 | 3.6 ±1.3 | 0.994±0.003 |

Values are presented as mean ± SE. Differences statistically not significant ( $P > 0.05$ ), Mann Whitney test. BIOM, Biological Observation Matrix; FEP, fibroepithelial polyp; OSCC, oral squamous cell carcinoma.

### 3.5 Overall mycobiome compositional profile

A total of 364 species belonging to 162 genera and two phyla were detected in the samples. The number of species per sample ranged from 4 to 64. However, only 74 genera and 125 species were identified in more than one sample: seven genera and 10species in ≥25% of the samples, and four genera and five species in >50%.

### 3.6 Fungal species richness and diversity of OSCC versus FEP

The number of species per sample ranged from 4 to 29 for the cases and from 8 to 64 for the controls. The FEP controls had significantly higher species richness and α-diversity than the cases (Table 2). Rarefaction curves show that as few as 1,500 reads per sample represented sufficient sequencing depth (Figure 2a). No separate clusters formed for the cases and controls by PCoA (Figure 2b).

**Table 2:** Species richness, α-diversity and coverage (mean±SE) calculated from the rarefiedbiom.

| Group | Observed richness* | Chao1* | Shannon index* | Good's coverage |
| --- | --- | --- | --- | --- |
| OSCC | 11.7±6.4 | 13.6±7.0 | 1.5±0.9 | 0.999±0.000 |
| FEP | 17.7±9.8 | 19.9±12.2 | 2.1 ±1.2 | 0.999±0.001 |

\* $p \leq 0.05$ (Mann–Whitney test). OSCC, oral squamous-cell carcinoma; FEP, fibro-epithelial polyps.

**Figure 2:**
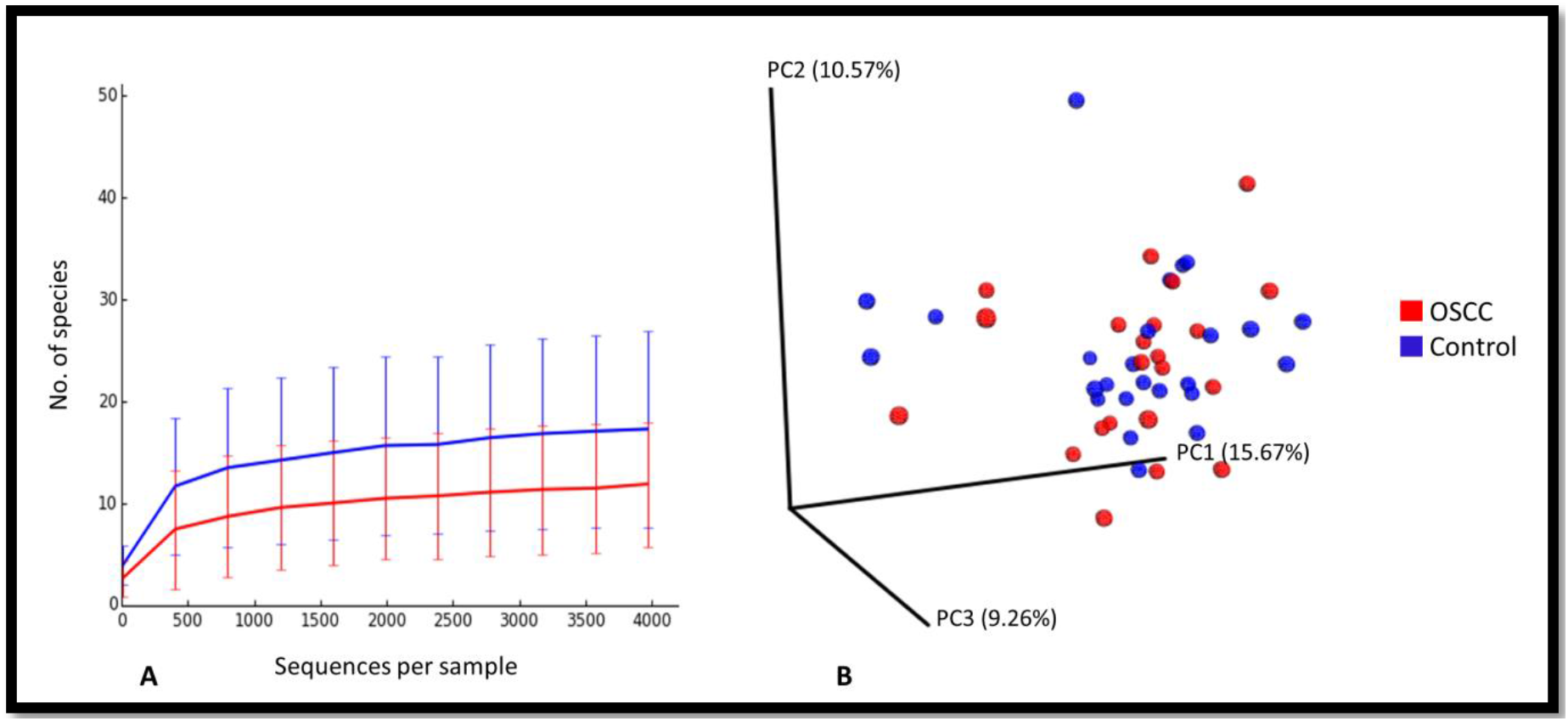
Rarefaction and β-diversity. (a) Rarefaction curves showing the number of observed species as a function of sequencing depth. (b) Non-clustering of the study subjects by principal components analysis (weighted-Unifrac).

### 3.7 Bacterial dysbiosis by differentially abundant taxa

The differentially abundant genera and species between the cases and controls are delineated in Figure 3. Genera *Capnocytophaga, Pseudomonas*, and *Atopobium* were associated with OSCC, while *Lautropia, Staphylococcus, Propionibacterium*, and *Sphingomonas* were the most significantly abundant in FEP. At the species level, *Campylobacter concisus, Prevotella salivae, Prevotella loeschii*, and *Fusobacterium oral taxon 204* were abundant in the cases, while 7 *Streptococcus* species, 2 *Rothia* species, *Lautropia mirabilis*, and *Leptotrichia* oral taxon 225, among others, were significantly enriched in the controls.

**Figure 3.**
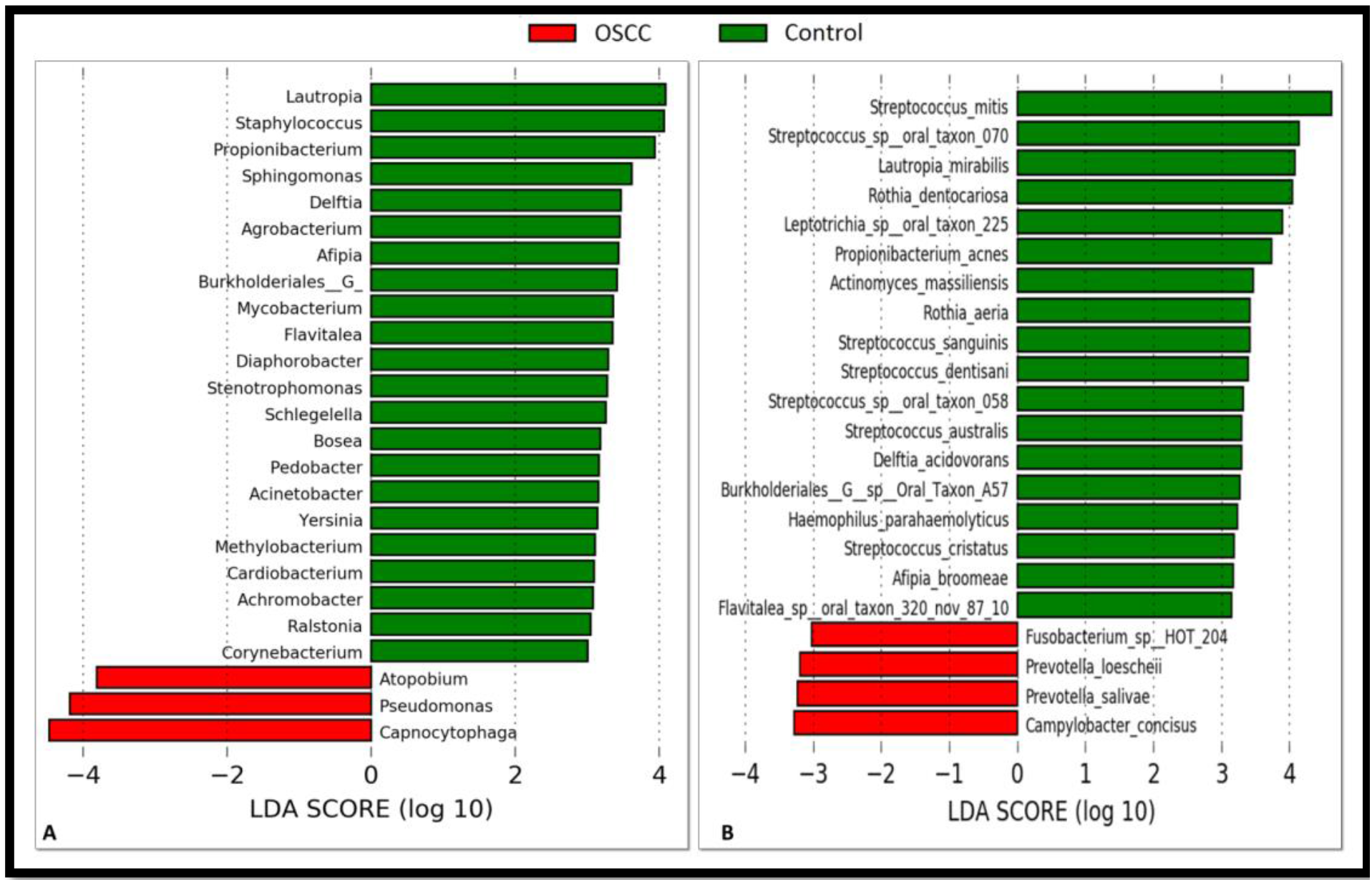
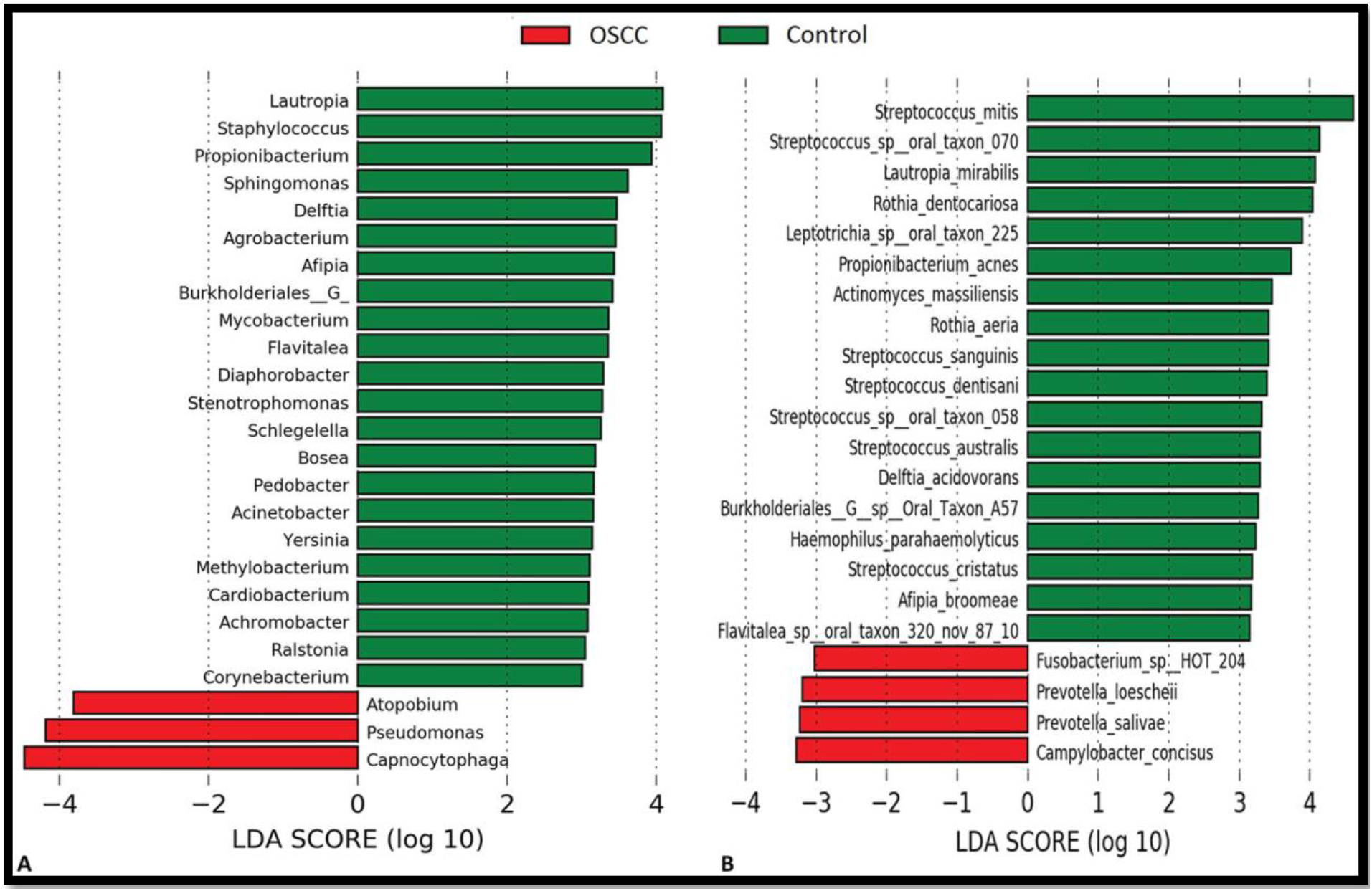
Differentially abundant taxa. (**A**) Genera and (**B**) species that were significantly more abundant between the cases and controls as identified by linear discriminant analysis (LDA) effect size analysis: linear discriminant analysis score ≥3. OSCC, oral squamous cell carcinoma

### 3.8 Fungal dysbiosis by differentially abundant taxa

The differentially abundant genera and species in cases and controls were detected by LEfSe (Figure 4). The genera *Candida, Hannaella*, and *Gibberella* were significantly abundant in OSCC. On the contrary, *Trametes* and *Alternaria* were lavishly present in FEP. When it comes to species level, *C. albicans, C. etchellii*, and a potentially novel species phylogenetically related to *Hannaella luteola* were significantly plentiful in OSCC. Although, *C. albicans* was identified in 100% of the samples, the average relative abundance in OSCC was twofold higher than that in the controls (61.2% vs. 29.6%). *C. etchellsii* was identified in 32% of the cases and 8% of the controls. On one hand, *Hannaella luteola*–like species was detected in 20% of OSCC samples but none of the controls. On the other hand, a potentially novel *Hanseniaspora uvarum*–like species, in addition to *M. restricta, A. tamarii, Cladosporium halotolerans, Alternaria alternata*, and *Malassezia furfur* were exclusively found in FEP.

### 3.9 Oral risk habits of OSCC cases and FEP controls of dysbiotic mycobiome

As shown in table 3, betel quid chewing, smoking and alcohol consumption were the risk habits of this study group. Among cases the mean duration of betel chewing was 34.09 years (SD±12.87) and the mean number of betel quids chewed per day was 14.65 (SD±7.30). In contrast, the mean duration of betel chewing of controls was 17.92 years (SD±15.34) and the mean number of betel quids chewed per day was 3.88 (SD±3.65), much less than that of cases. The mean duration of smoking among cases was 24.32 years (SD±15.45), whereas this duration was 13.76 years (SD±13.85), in controls. In addition, the mean duration of alcohol consumption was 29.45 years (SD±10.43) in cases and 17.20 years (SD±12.46) in controls. In overall, for all habits the mean durations were longer for Cases compared with Controls and those differences were statistically significant (p<0.05).

**Table 3:** Distribution of the Cases and Controls by well-established risk factors: Betel Chewing Practice, Smoking & Alcohol Consumption.

| Variable | Cases n=22 | Controls n=25 | p- value |
| --- | --- | --- | --- |
| Mean ± SD duration of Betel chewing in years! | 34.09± 12.87 | 17.92±15.34 | 0.001(p<0.05)** |
| Mean ± SD number of betel quids chewed per day! | 14.65± 7.30 | 3.88± 3.65 | 0.001(p<0.05)** |
| Mean ± SD duration of Smoking in years! | 24.32± 15.45 | 13.76±13.85 | 0.017(p<0.05)** |
| Mean ± SD duration of alcohol consumption in years! | 29.45± 10.43 | 17.20± 12.46 | 0.001(p<0.05)** |
| **Type of Betel Chewing Habit \*** | **N %** | **N %** | **p- value** |
| Never | 0 (0.0) | 4(16.0) | 0.021(p<0.05)** |
| Past | 3(13.6) | 2(32.0) |  |
| Sometimes | 1( 4.6) | 7(28.0) |  |
| Daily | 18 (81.9) | 12(48.0) |  |
| **Total** | **22(100.0)** | **25 (100.0)** |  |
| **Type of Smoking Habit \*** | **N %** | **N %** | **p-value** |
| Never | 4(18.2) | 8(32.0) | 0.555(p>0.05) |
| Past | 5(22.7) | 7(28.0) |  |
| Sometimes | 4(18.2) | 4(16.0) |  |
| Daily | 9(40.9) | 6(24.0) |  |
| **Total** | **22(100.0)** | **25 (100.0)** |  |
| **Type of Alcohol Consumption Habit \*** | **N %** | **N %** | **p-value** |
| Never | 0(0.0) | 4(16.0) | 0.001(p<0.05)** |
| Past | 5(22.7) | 2(32.0) |  |
| Sometimes | 4(18.2) | 15(60.0) |  |
| Daily | 13(59.1) | 4(16.0) |  |
| **Total** | **22(100.0)** | **25 (100.0)** |  |

t-test for comparison of means of independent samples +
Chi Square Test of Statistical Significance,\*

Furthermore, as revealed by Table 3, the overwhelming majority, (91.8%) of case were current betel chewers, who consumed betel quids on a daily-basis. In contrast, 48.0% of controls consumed betel quids on daily basis. However, 4.6% of cases and 28.0% of controls consumed betel quid, some-times but not on a daily basis. Furthermore, past chewers were reported to be 13.6% and 32.0% for the cases and controls respectively. Nevertheless, all cases had past or present betel chewing habit and none were never chewers.. In contrast, 16% of controls were “never chewers’ of betel quid. As shown in Table 3, a higher proportion of Cases were daily betel chewers compared to Controls and this difference was statistically significant (p<0.05).

According to the present classification of type of smoking habit, almost 41.0% of cases were present smokers who got used to practice this habit on a daily basis. In contrast,, 24.0% of controls smoked daily. Moreover, 18.2% of cases and 16.0% controls were present smokers who smoked sometimes but not on daily basis. On the other hand, never smokers were 18.2% and 32.0% for the cases and controls respectively. Thus, as shown in Table 3, higher proportion of Cases were daily smokers compared to Controls, however this difference was not statistically significant (p>0.05).

As per Table 3, nearly 59.0 % of cases were *current drinkers*, who consumed alcohol daily but only 16.0% of controls were addicted to this habit on a daily basis. Furthermore, 18.2% of cases and 60.0% of controls were drinking alcohol sometimes. Past drinkers were reported as 22.7% and 28.0% for the cases and controls respectively. There were 0.0% of *never* alcohol users among cases, and 16.0% of *never users* among controls. Thus, higher proportion of Cases were daily drinkers compared to Controls and this difference was statistically significant (p<0.05).

## 4. Discussion

The present study corroborates the notion that microbial dysbiosis may be associated with the progression of oral carcinogenesis via chronic inflammation [6, 13] and agree with the findings of inflammatory bacteriome and OSCC [9], A dysbiotic mycobiome dominated by *Candida albicans* is identified within oral squamous-cell carcinomas [10] and other metagenomic epidemiological studies which found associations of dysbiotic microbiome with OSCC progression [14,15] published previously. Furthermore, the present study reminded us that microbial dysbiosis is closely related to the aetiopathogenesis of colon, gastric, esophageal, pancreatic, breast, and gall bladder carcinomas [9, 16], modulating the tumor microenvironment by virulence factors, metabolites, toxins [6, 13] and immune evasion of pathobiont essentially favorable to neoplastic cells to triumphant with immune checkpoints, immune editing and avoid immune elimination [17].

Alpha (ἀ) diversity is a summary statistic to understand the species richness of a meta-community in a clinical or environmental sample at a given time [4]. It is important to note that the Sri Lankan oral squamous cell carcinoma (OSCC) tissue samples demonstrated lower bacterial species richness and diversity (see Figure 1) compared to the FEP controls [9]. Microbial dysbiosis and immunosuppression might explain this observation, although the alpha diversity was statistically insignificant [9]. This finding contradicts previous studies that used saliva or swabs rather than properly sterilized tissue samples or contra lateral samples of the same patient as controls not considering the *field characterization* even in clinically normal sites [18, 19]. It is essential to highlight that the FEP controls exhibited statistically significantly higher species richness and alpha diversity [9] (see Table 2) compared to the cases in a study conducted by the same authors of the present research, which identified a dysbiotic mycobiome predominantly composed of *Candida albicans* within oral squamous cell carcinoma [10].

Beta (β) diversity is a similarity index to compare two groups. In the present study, obviously the dysbiotic bacteriome separated into two clusters as per cases and controls by principal components analysis (PCoA) (weighted Unifrac) (Figure 1). This discovery is consistence with the finding of OSCC specific bacteriome in tumour micro environment [4, 7, 9, 15, 19, 20]. In contrast, non-clustering of cases and controls separately was observed in the mycobiome study [10] due to inherent limitations as described rationally [10] of exploring the mycobiome compared the bacteriome [9].

The differentially abundant genera (except Atopobium) [9] and species stated in the result section appeared to be associated with oral cancer in previous close-ended as well as open-ended molecular studies to unveil the composition profile of bacteriome of OSCC by other authors [7,15, 19,20]. We found a significant association of *Atopobium* for the first time [9]. The handful taxa *Capnocytophaga, Pseudomonas, Atopobium* and *Campylobacter concisus, Prevotella salivae, Prevotella loeschii*, and *Fusobacterium oral taxon 204* were able to flourish as differently abundant taxa, enriched in the OSCC tumor microenvironment as described previously are putative periodontopathogens. Interestingly, *Pseudomonas* was associated with periodontitis but others were putative periodontal pathogens thus, aetiological agents of chronic periodontitis of periodontal disease [7, 9, 14, 21]. Candida, namely *C. albicans* and *C. etchellsii*, were significantly differentially abundant in OSCC after applying high-flown LEfSe in this study and another study published by the same authors [10]. Other than the well-documented facts that candidiasis triggers type 3 responses initiated by activated IL-17-secreting cells [22] production of carcinogenic levels of acetaldehyde by *C. albicans* and few other *Candida* species as well as high potential of some *C. albicans* strains to produce nitrosamines have been experimentally shown to induce dysplasia [10], the causative role of Candida in oral carcinogenesis yet to be proven [10, 22]. Our findings are on par with the *Candidal blooms* detected in OSCC patients in an increasing number of prevalence and epidemiological studies [10, 22].

In the present study, OSCC patients were also heavier users of betel-quids with smokeless tobacco and areca nut, cigarettes, and alcohol with the duration of substance abuse (p<0.05)**. Hence, there was a statistically significant difference in betel-quid chewing and alcohol consumption habits among OSCC cases and FEP controls. These oral risk habits may introduce inflammogenic, mutagenic, and carcinogenic substances to the oral epithelium, and, the duration of substance abuse is directly proportional to having orally potential malignant disorders (OPMD) and oral cancer (OSCC) s [23]. Furthermore, these factors consider environmental factors inducing selection pressure [22,23,24] on indigenous microflora to elevate patho-bionts disproportionately who are putative periodontal pathogens [18,21,22,23,24] suppressing commensals mainly due to the overburdened immune functions in tumor microenvironment and inflamed oral mucosa of substance abusers to keep them under control. Chronic inflammation or persistent inflammation seems conducive to neoplastic transformations [6]. Findings of the present study and a few publications based on pioneering work investigating compositional and functional aspects of metagenome of a group of Sri Lankan male OSCC patients paved the way to generate similar findings subsequently based on epidemiological studies using NGS technologies [14, 25]. The role of microbial dysbiosis as a potential cause of oral carcinogenesis remains unproven, and the in vitro experiments conducted so far have yielded inconclusive results in this area. To date, no animal models have been able to induce oral carcinogenesis through a mono or poly periodontal pathogens. However, emerging evidence suggests that interactions between periodontal pathogens and neoplastic cells, as well as immune cells may contribute to the progression of carcinogenesis in a few animal models, [6, 14, 18]. At the same time, polymicrobial interactions may result in bioactive, antimicrobial and anti-inflammatory crude compounds to suppress the competitors as these microbes compete with each other for attachment sites and nutrients even in the OSCC tumor microenvironment. Isolation, purification, and identification of these anti-cancers compounds an emerging research arena and seem a subspecialty of natural product chemistry [10, 13].

The study design was mainly due to the diagnosis of FEP at a younger age than the cases, a significant age difference between the cases and control, the small size of the representative subsample, the impossibility of matching the controls for oral risk habits or adjusting for their confounding effects statistically were inherent limitations of this unmatched case-control study. The presence of PCR inhibitors, absence of the optimum surface sterilization to avoid saliva contamination 100%, not optimizing the DNA extraction protocol to a maximum recovery of fungal DNA, exaggeration in species identification with hundreds of novel species, and other inherent limitations in metagenomic studies are also cannot be forgotten.

## 5. Conclusion

Our study suggests a few potential periodontal pathogens, including *Campylobacter concisus, Prevotella salivae, Prevotella loeschii*, and *Fusobacterium oral taxon 204*, as well as *Candida etchellii*. These eligible candidates could be used in mechanistic studies to explore their roles either in the initiation or progression of oral squamous cell carcinoma (OSCC), similar to previous research involving *Porphyromonas gingivalis* and *Fusobacterium nucleatum*. Additionally, to investigate Candida albicans further to understand the signaling mechanisms that may contribute to oral carcinogenesis. It is also important to identify the most promising species from the *Capnocytophaga* and *Atopobium* genera for use in both in vivo and in vitro studies. Furthermore, betel quid chewing and alcohol consumption have been shown to have a statistically significant association with dysbiosis in the tumor microenvironment.

## 6. Recommendations and future directions

Future work on microbial ecological *metagenomics, metatranscriptomics, metaproteomics*, and *metabolomics* with cohort study design (including adequate sample size, controlling for confounding effects of established risk factors) could be recommended as a re-emerging arena of groundbreaking research into cancers in general and for oral cancer, in particular, to revolutionize the landscape of time-varying microbiome (microbial dysbiosis) in the natural history of carcinogenesis, to identify key microbiota or shifts in community structure to determine whether the microbiota are *cause* or *effect* of cancer risk due to favorable tumor microenvironment for flourishing certain microbes, their genetic potential, their proteins and metabolites.

The *one health approach* and revelation of evolution and dissemination of *gut and environmental resistomes* as well as the virulence of multi-drug resistant healthcare-associated infectious agents reiterate the importance of revisiting *microbiome-first-medicine considering* the human as a *supra organism* not neglecting the indigenous flora specific to each individual in healthy and disease conditions govern by the social determinants, sociodemographic profile and health disparities unfavourable to the most vulnerable. There is a need to incorporate the diagnostic biomarkers and risk panels in routine clinical laboratory standard operating procedures (SOP) s at least in national and regional reference laboratories to supplement the clinical utility and reproducibility as the efficacy and usefulness of histopathology in predicting malignant transformation of OPMD even in the absence of oral epithelial dysplasia (OED). As there is promising evidence in ‘omics’ studies it is worth validating the diagnostic biomarkers or the microbes produce metabolites that may increase the risk of carcinogenesis as early detection prevent cancer associated deaths. Furthermore, prognostic biomarkers or the microbiota produce anti-cancer, antimicrobial, and anti-inflammatory substances to control cancer-related death by improving the 5yrs survival rates.

## Acknowledgments

We acknowledge late Professor Newell Johnson, Associate Professor Glen Ulett, Professor of Microbiology, School of Medical Science and Pharmacy, Gold Coast Campus, Griffith University, QLD 4222, Prof. Nezar Al-Hebshi, Professor in Dental Research and Oral Microbiome, Maurice H. Kornberg School of Dentistry, Temple University, Philadelphia, USA, Dr. Tsute Chen, Associate Investigator Department of Oral Medicine, Infection and Immunity, Harvard School of Dental Medicine, Dr. Deepak Ipe and Dr. DJ Speicher for their valuable contribution to make this study success. We extended our gratitude to Prof. WM Tilakaratne, Senior Professor of Oral Pathology, and Prof. L. Samaranayake Professor of Oral Microbiology for their guidance. We thank Oral and Maxillo-Facial Surgeons Dr. Sharika Gunathilake, Dr. S.A.K.J. Kumara, Dr. Ranjith Lal Kandewatte, Dr. P. Kirupakaran, Dr. D.K. Dias, Dr. Chamara Athukorale, Dr. Suresh Shanmuganathan, and Dr. T. Sabesan for facilitating data and sample collection from their respective Oral and Maxillo-Facial Units.

